# ProbeTools: Designing hybridization probes for targeted genomic sequencing of diverse and hypervariable viral taxa

**DOI:** 10.1101/2022.02.24.481870

**Authors:** Kevin S. Kuchinski, Jun Duan, Chelsea Himsworth, William Hsiao, Natalie A. Prystajecky

**Affiliations:** Department of Pathology and Laboratory Medicine; University of British Columbia; Vancouver, British Columbia, Canada; Animal Health Centre; British Columbia Ministry of Agriculture, Food, and Fisheries; Abbotsford, British Columbia, Canada; School of Population and Public Health; University of British Columbia; Vancouver, British Columbia, Canada; Faculty of Health Sciences; Simon Fraser University; Burnaby, British Columbia, Canada; Public Health Laboratory; British Columbia Centre for Disease Control; Vancouver, British Columbia, Canada

**Keywords:** Influenza A viruses, avian influenza viruses, viral genomics, hybridization probe capture, targeted genomic sequencing, viral surveillance

## Abstract

**Background:** Sequencing viruses in many specimens is hindered by excessive background material from hosts, microbiota, and environmental organisms. Consequently, enrichment of target genomic material is necessary for practical high-throughput viral genome sequencing. Hybridization probes are widely used for enrichment in many fields, but their application to viral sequencing faces a major obstacle: it is difficult to design panels of probe oligo sequences that broadly target many viral taxa due to their rapid evolution, extensive diversity, and genetic hypervariability. To address this challenge, we created ProbeTools, a package of bioinformatic tools for generating effective viral capture panels, and for assessing coverage of target sequences by probe panel designs *in silico*. In this study, we validated ProbeTools by designing a panel of 3,600 probes for subtyping the hypervariable haemagglutinin (HA) and neuraminidase (NA) genome segments of avian-origin influenza A viruses (AIVs). Using *in silico* assessment of AIV reference sequences and *in vitro* capture on egg-cultured viral isolates, we demonstrated effective performance by our custom AIV panel and ProbeTools’ suitability for challenging viral probe design applications.

**Results:** Based on ProbeTool’s *in silico* analysis, our panel provided broadly inclusive coverage of 14,772 HA and 11,967 NA reference sequences. 90% of these HA and NA references sequences had 90.8% and 95.1% of their nucleotide positions covered *in silico* by the panel respectively. We also observed effective *in vitro* capture on a representative collection of 23 egg-cultured AIVs that included isolates from wild birds, poultry, and humans and representatives from all HA and NA subtypes. 42 of 46 HA and NA segments had over 98.3% of their nucleotide positions significantly enriched by our custom panel. These *in vitro* results were further used to validate ProbeTools’ *in silico* coverage assessment algorithm; 89.2% of *in silico* predictions were concordant with *in vitro* results.

**Conclusions:** ProbeTools generated an effective panel for subtyping AIVs that can be deployed for genomic surveillance, outbreak prevention, and pandemic preparedness. Effective probe design against hypervariable AIV targets also validated ProbeTools’ design and coverage assessment algorithms, demonstrating their suitability for other challenging viral capture applications.

## BACKGROUND

Most viral specimens are characterized by low amounts of viral genomic material and excessive background from viral hosts and environmental organisms. Consequently, practical viral genome sequencing requires targeted enrichment for confident detection and accurate genotyping, especially in high-throughput surveillance and clinical applications [1-3]. Hybridization probe capture methods have been used for viral target enrichment [4-7], but designing probe oligo sequences for many viruses can be a major obstacle due to extensive genomic diversity and hypervariability within and between viral taxa [8-13].

Probe panels are typically designed by enumerating probe-length sub-sequences (k-mers) from reference sequences. The simplest approach to designing probes for hypervariable taxa is to enumerate k-mers from an exhaustive collection of reference sequences, thereby including as much genomic divergence in the design space as possible [7-8]. This approach becomes problematic, however, when redundant probe sequences are enumerated from repeated instances of conserved genomic loci.

A few strategies have been used to address this redundancy problem. One common strategy is to cluster similar k-mers after they have been enumerated [6-7]. Another strategy is to align candidate probe sequences against select reference genomes to identify and retain only those probes targeting divergent genotypes [8]. Redundancy has also been addressed by constraining the design space to a limited number of representative reference genomes, selected either by manual curation or clustering [9-12]. Some of these strategies have been supplemented with multiple sequence alignments over hypervariable loci or entire genomes so that probes are designed from consensus and degenerate sequences [9-10].

Spacing between probe sequences is another complicated design consideration. Regular spacing (tiling) is the most common approach because it is easy to implement, but it does not ensure optimal positioning of probes. Reducing the spacing increases the likelihood that some enumerated probes are optimally positioned, but it also increases the number of probe candidates and any associated computation to collapse redundancy among them. Creating the smallest possible panel of probes that optimally covers the entire target space quickly becomes an intractable computational problem, one that had led to increasingly complicated approaches including sophisticated minimization of loss functions [13].

Efforts to address viral hypervariability have resulted in several elaborate probe design algorithms. Unfortunately, these have largely been implemented on a study-by-study basis and have not resulted in general-purpose software tools that can be easily used by others. Meanwhile, among the handful of published probe design packages, there is only one option that specifically addresses viral hypervariability [13]. The rest are intended for comparatively conserved eukaryotic genomes and are inadequate for many viral applications [14-17]. This leaves virologists with limited options for designing their own hybridization probes, especially if they have minimal capacity for custom programming, sophisticated mathematics, and experimental bioinformatics.

Here, we present ProbeTools, a user-friendly command line software package for designing compact probe panels against diverse viral taxa and other hypervariable genomic targets. It provides easy-to-use modules for generating probes and assessing panel coverage of provided target sequences. We demonstrate ProbeTools’ effectiveness by designing capture panels for avian-origin influenza A viruses (AIVs). These viruses are subtyped by two hypervariable viral surface proteins called haemagglutinin (HA) and neuraminidase (NA), making them an appropriately challenging case study for ProbeTools. The genome segments encoding these proteins have diversified into 16 avian-origin HA subtypes and 9 avian-origin NA subtypes, giving rise to 144 possible combinations and the HxNx nomenclature used in both animal and human contexts (*e*.*g*. H1N1, H3N2, H5N1, H7N9). Furthermore, each of these subtypes has diverged into numerous clades, many of which have been only partially characterized [12, 18-19].

AIV lineages have varying potential for spillover from wild birds into poultry and humans [20-25], posing a perennial threat to agriculture and public health. Some lineages cause costly outbreaks of severe disease in poultry flocks which, in turn, expose humans to potentially dangerous zoonotic influenza infections. This threatens economic disruption, future pandemic crises, and new types of seasonal influenza, which remains an important global health burden and among the ten leading causes of death worldwide [12, 21-31]. Consequently, surveillance of AIVs in wild birds is a cornerstone of outbreak prevention and pandemic preparedness [12, 20, 32-33]. An effective panel of AIV-specific probes would be instrumental for these genomics-based surveillance efforts.

In this study, we designed and validated a compact panel of 3,600 probes for detecting and subtyping AIVs. Our results showed broad inclusivity against all avian-origin HA and NA subtypes based on *in silico* predictions against of tens-of-thousands of AIV reference sequences. We also demonstrated successful captures *in vitro* on a representative collection of 23 egg-cultured AIVs.

## RESULTS

### Assessing basic k-mer clustering and marginal improvements to target coverage with additional probes

We began by assessing probe design against hypervariable targets with a basic k-mer clustering algorithm, wherein all 120-mers were enumerated from a target space of AIV reference sequences then clustered based on 90% nucleotide sequence identity. We used this strategy, implemented in the ProbeTools *clusterkmers* module, to generate probe panels of increasing size against 14,772 HA segment reference sequences and 11,967 NA segment reference sequences. We then used the ProbeTools *capture* module, which aligns probe sequences against target sequences, to assess target space coverage, *i*.*e*. the percentage of nucleotide positions in each target sequence covered by at least one probe in the panel (Figure 1A, solid lines). As expected, panels with more probe sequences provided better target space coverage, however we observed diminishing marginal improvements for both HA and NA genome segments. We also noted that reference sequences with no probe coverage remained in the target space past the point of diminishing marginal returns. These results highlighted two limitations of the basic k-mer clustering approach: HA and NA segments remained undetected despite designing additional probes, and additional probes provided only modest and diminishing improvements to the distribution of target coverage.

**Figure 1:**
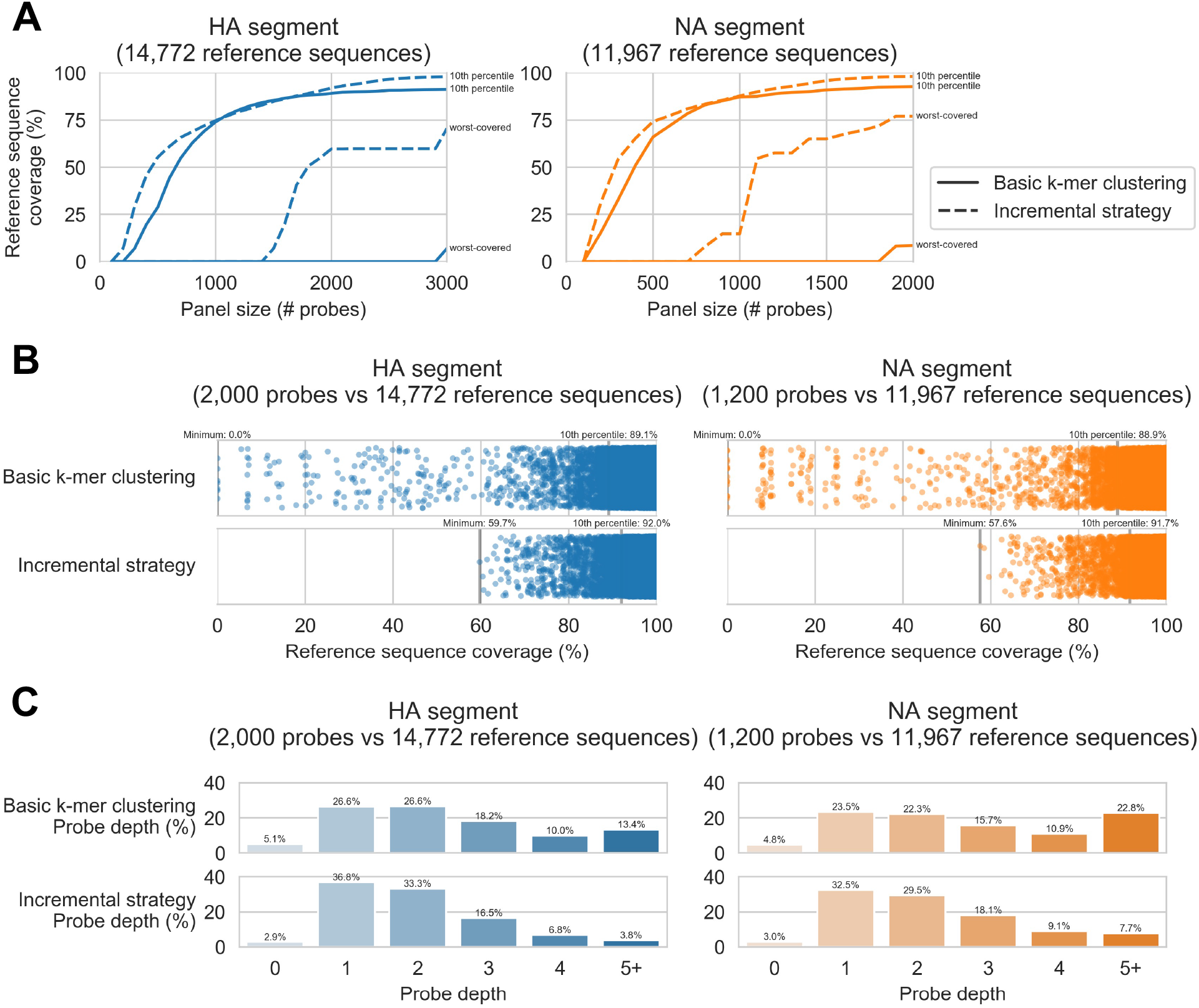
Incremental design strategy improves upon basic k-mer clustering for probe panel design. Panels were designed against target spaces of 14,772 haemagglutinin (HA) and 11,967 neuraminidase (NA) genome segment reference sequences. The ProbeTools *clusterkmers* module was used to make panels using basic k-mer clustering and the *makeprobes* module was used to make panels with an incremental strategy. For each panel, probe coverage of reference sequences was assessed *in silico* using the ProbeTools *capture* module. A) For both strategies, increasing panel size improved the 10^th^ percentile of reference sequence coverage with diminishing marginal increases, but incrementally designed panels achieved more extensive coverage at larger panel sizes. Incrementally designed panels also provided better coverage of the worst-covered reference sequence using fewer probes. B) Incrementally designed panels shifted coverage distributions upwards for the worst-covered reference sequences. Each reference sequence in the target space is represented as a dot, plotted according to its probe coverage. Coverage of the worst-covered reference sequence and 10^th^ percentile of all reference sequences are indicated above the axis. C) Incrementally designed panels improved reference sequence coverage by re-distributing probes from regions with deep coverage (4 or more probes) to regions with shallow coverage (2 or fewer probes).

### Improving target coverage with incremental panel design focused on poorly covered targets

To address the limitations we observed with basic k-mer clustering, we devised an incremental design strategy to improve marginal coverage increases, especially for poorly covered targets. In this strategy, basic k-mer clustering was used to design panels in smaller batches of 100 probes. After adding each batch to the growing panel, target space regions without probe coverage were identified using the *capture* module. These low coverage regions were then extracted with another ProbeTools module called *getlowcov* and used as a new target space for designing the next batch. In this way, each subsequent batch of probes was focused on regions not already covered by the panel.

We compared target space coverage for panels designed with this incremental strategy against panels designed above using basic k-mer clustering (Figure 1). The incremental strategy provided higher 10^th^ percentiles of coverage, especially for HA panels larger than 2000 probes and NA panels larger than 1200 probes (Figure 1A). Furthermore, the incremental strategy provided better coverage for the worst-covered reference sequences (Figure 1AB). We also compared depth of probe coverage, *i*.*e*. the number of probes covering each nucleotide position in target sequences (Figure 1C). This comparison suggested that the incremental strategy improved target coverage by redistributing probes from positions with deep coverage to shallow coverage. Based on the observed performance improvements of the incremental strategy, it was implemented as an additional self-contained ProbeTools module called *makeprobes* (Figure 2).

**Figure 2:**
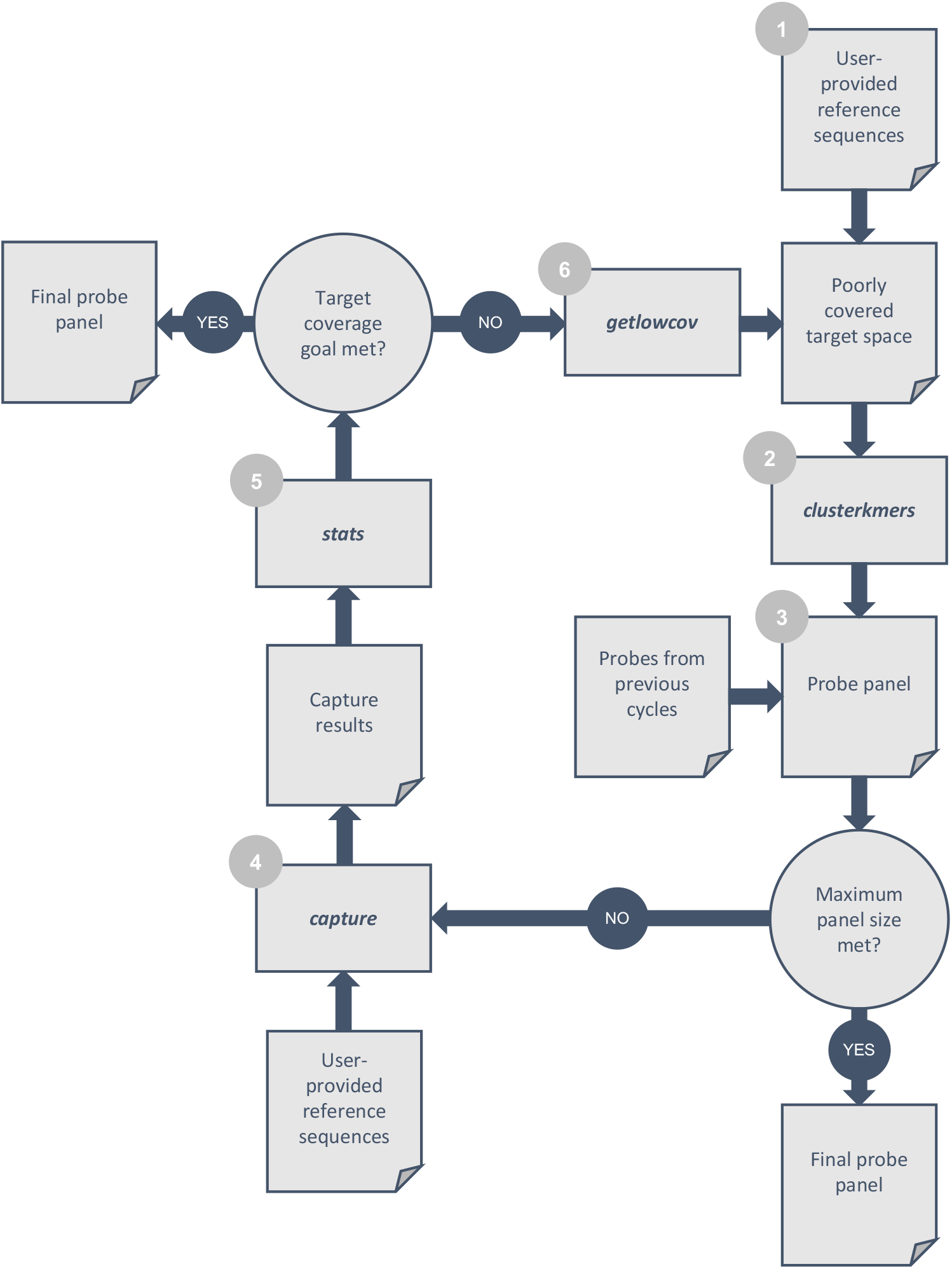
ProbeTools *makeprobes* module implements a generalized incremental design algorithm. 1) The user provides a FASTA formatted file containing target sequences, which forms the total target space and become the poorly covered target space for the first loop of the design cycle. 2) The ProbeTools *clusterkmers* module generates a batch of probe sequences from the poorly covered target space using its k-mer clustering algorithm. 3) The latest batch of probes is combined with probes from previous batches to generate the current probe panel. If the size of the current probe panel meets the maximum panel size set by the user, the design loop ends and the current panel becomes the final panel, otherwise… 4) The ProbeTools *capture* module determines which nucleotide positions in the total target space are covered by the current probe panel. 5) The ProbeTools *stats* module calculates the 10^th^ percentile of target coverage from the *capture* module results. If the target coverage goal set by the user is met, the current probe panel becomes the final probe panel, otherwise… 6) The *getlowcov* module extracts low coverage regions of the target space from the *capture* module results. These become the new poorly covered target space, and the design loop repeats.

### Predicted coverage of HA and NA subtypes by AIV_v1 panel

Using the incremental strategy implemented in the ProbeTools *makeprobes* module, we generated an AIV-targeting probe panel called AIV_v1. It was composed of 1,935 HA-specific probes and 1,435 NA-specific probes. We also included 184 probes targeting the highly conserved matrix segment (M) which is the standard AIV diagnostic target [24, 38]. We then used the ProbeTools *capture* module to predict probe coverage using the AIV_v1 panel for all 36,313 AIV reference sequences in the target space. The minimum, maximum, and 10^th^ percentile of reference sequence coverage was calculated for each HA and NA subtype and the M segment (Figure 3A).

**Figure 3:**
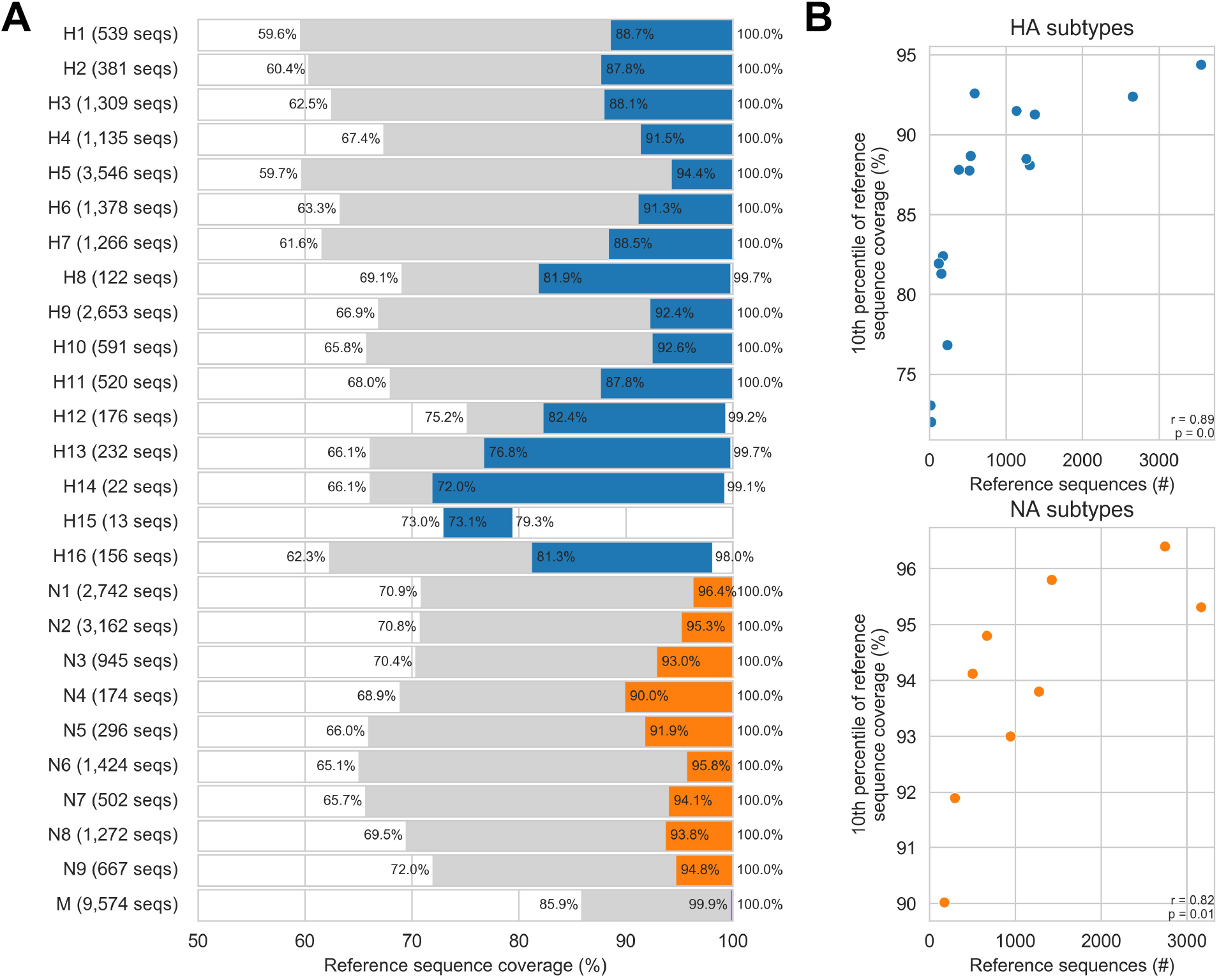
The ProbeTools-designed AIV_v1 panel provided broadly inclusive coverage *in silico* of avian-origin HA subtypes, NA subtypes, and M segments. The AIV_v1 panel of 3,600 probes was designed using the ProbeTools *makeprobes* module. It was composed of 1,935 haemagglutinin (HA) segment-specific, 1,435 neuraminidase (NA) segment-specific, and 184 matrix (M) segment-specific probes. A) Coverage predictions against 36,313 reference sequences were generated with the ProbeTools *capture* module and stratified by subtype for HA and NA segments. The minimum, 10^th^ percentile, and maximum of probe coverage against reference sequences from each subtype/segment are indicated. B) A significant positive monotonic association was observed between the number of sequences from a subtype in the target space and that subtype’s 10^th^ percentile of coverage. Each dot represents an HA or NA subtype, and the results of Spearman’s rank correlation test are indicated on the plots.

We observed that M segments had the best coverage followed by NA subtypes then HA subtypes, reflecting the comparative levels of genomic diversity within these genome segments. No reference sequence had less than 59.6% coverage, which is sufficient for segment and subtype identification. HA subtypes H5, H7, and H9 are considered high priority for AIV surveillance because they frequently cause agricultural outbreaks and novel influenza infections in humans [23-26, 38]; 90% of H5, H7, and H9 reference sequences had at least 94.4%, 88.5%, and 92.4% probe coverage respectively. We also noted a significant positive monotonic association between a subtype’s target coverage distribution and number of reference sequences from that subtype in the target space (Figure 3B). This indicated that over-representing subtypes in the target space resulted in preferential design and better probe coverage for these targets, *e*.*g*. the high priority subtypes H5, H7, and H9.

### *In vitro* capture of diverse egg-cultured influenza isolates

After assessing the AIV_v1 panel *in silico*, we had it synthesized and used it to perform *in vitro* captures on a collection of diverse egg-cultured AIV isolates (Table 1). We ensured that each avian-origin HA and NA subtype was represented in the collection, and we included isolates from wild birds, poultry, and humans. The collection contained 22 egg cultures, including one mixed infection, providing 23 viruses and 69 distinct HA, NA, and M segments for *in vitro* capture.

**Table 1:**
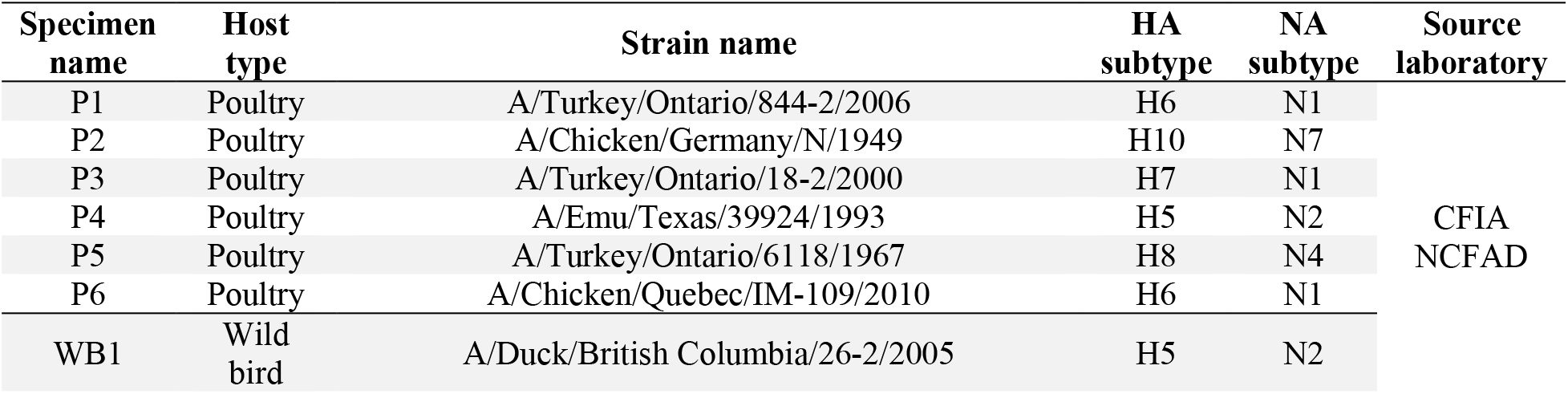
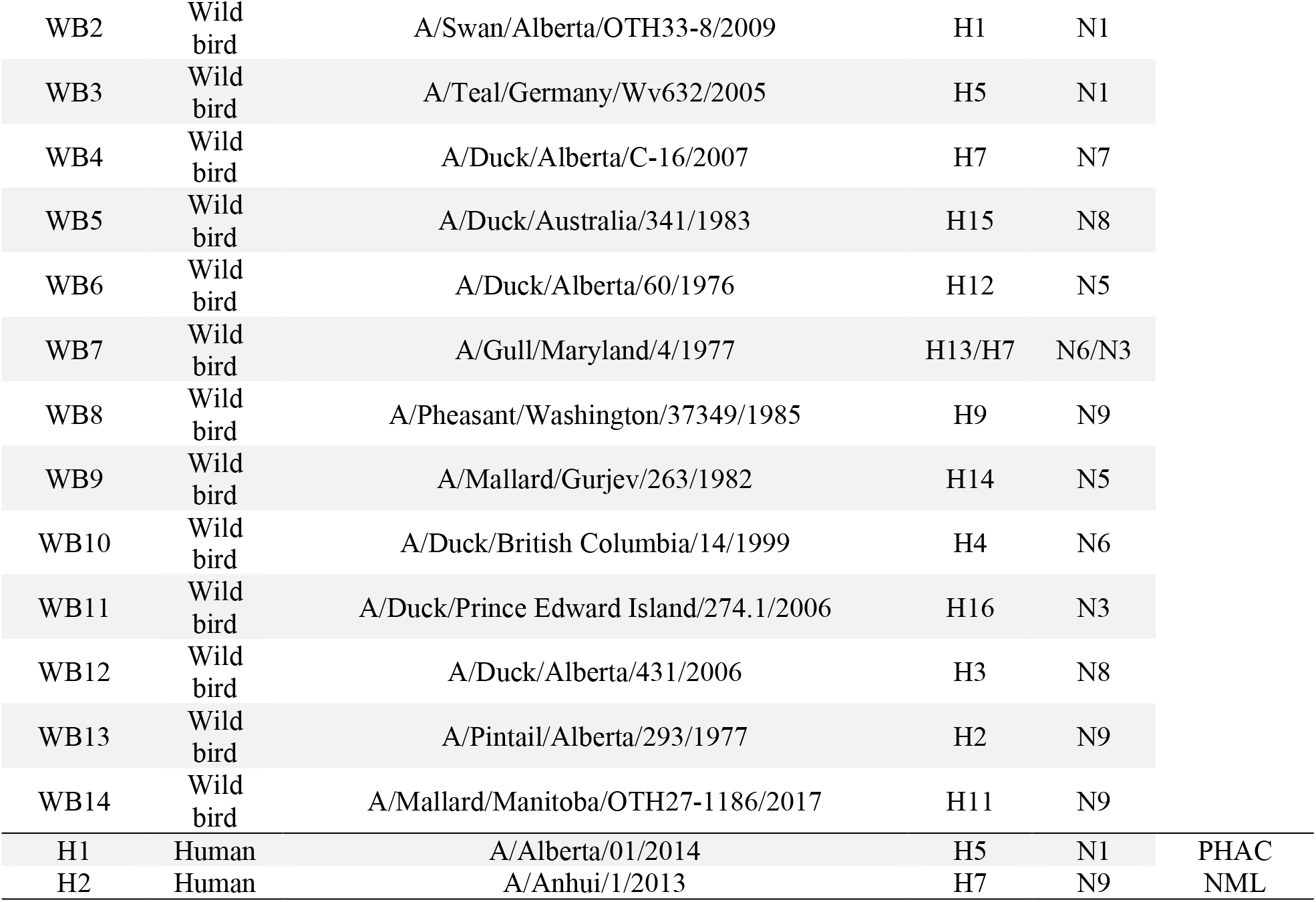
Representative collection of egg-cultured avian influenza virus isolates. Isolates were selected to provide representation of each avian-origin haemagglutinin (HA) and neuraminidase (NA) subtype as well as infections from poultry, wild bird, and human hosts. Each specimen was given a name based on an abbreviation of its host type and a sequential number (P for poultry, WB for wild bird, and H for human). Poultry and wild bird isolates were obtained from the Canadian Food Inspection Agency’s National Centre for Foreign Animal Disease (CFIA NCFAD), and human isolates were obtained from the Public Health Agency of Canada’s National Microbiology Laboratory (PHAC NML). Isolate subtypes were confirmed in-house by genome sequencing.

Sequencing libraries were prepared from each isolate then pooled. AIV library pools were diluted 1:100 (ng/ng) in libraries of background material made from mock-infected egg cultures, then captured three times independently using the AIV_v1 panel. Pre- and post-capture pools were sequenced to calculate mean fold-enrichment at each nucleotide position in these 69 HA, NA, and M segments. Half of all nucleotide positions had a mean fold-enrichment greater than 351.2-fold, and 90% of nucleotide positions had a mean fold-enrichment greater than 195.0-fold (Figure 4A). We also calculated the percentage of the capture pools composed of background material from the mock-infected egg cultures, then compared these percentages pre-and post-capture (Figure 4B). Before capture, the mean background percentage was 99.17%, but this was reduced to 0.03% following capture. Together, these data demonstrate effective enrichment of AIV material and removal of background by probe capture with the AIV_v1 panel.

**Figure 4:**
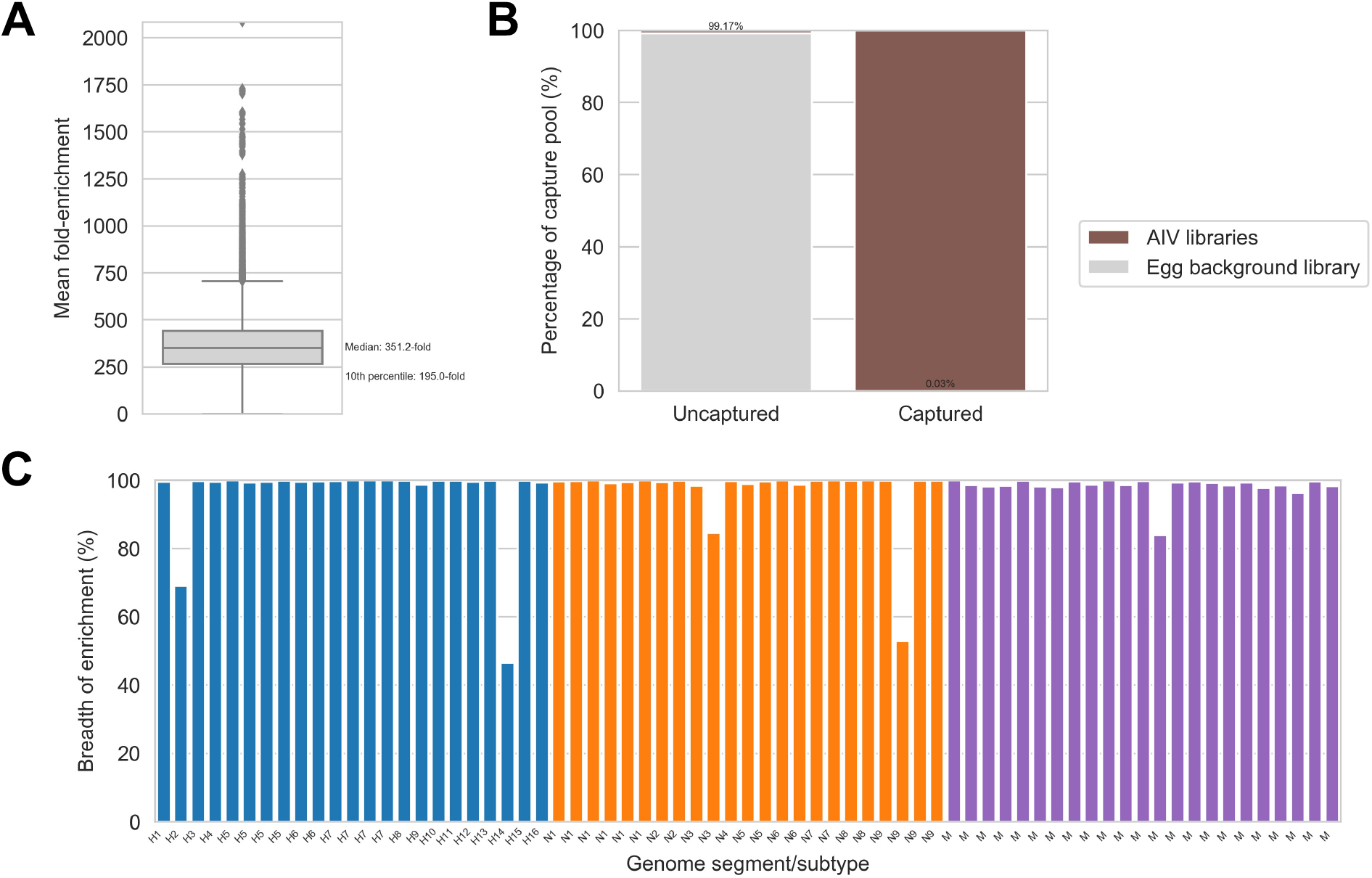
Effective *in vitro* capture of egg-cultured avian influenza virus isolates using the ProbeTools-designed AIV_v1 panel. The AIV_v1 panel of 3,600 probes was designed using ProbeTools, and it was used to capture sequencing libraries made from a representative collection of 23 egg-cultured avian influenza viruses (AIVs) (described in Table 1). AIV libraries were pooled together, diluted 1:100 (ng/ng) in libraries of background material made from mock-infected egg cultures, then captured three times independently. A) Pre- and post-capture pools were sequenced to calculate fold-enrichment at each nucleotide position in the haemagglutinin (HA), neuraminidase (NA), and matrix (M) genome segments of these isolates (mean of three independent replicates). B) Background material from mock-infected egg cultures was effectively removed during probe capture. C) Breadth of enrichment, *i*.*e*. the percentage of nucleotide positions that were significantly enriched by probe capture, was calculated for each HA, NA, and M genome segment in these isolates.

We also used these *in vitro* results to assess breadth of enrichment, *i*.*e*. the percentage of nucleotide positions in each HA, NA, and M segment that had been significantly enriched by probe capture (Figure 4C, Table S1). Breadth of enrichment was greater than 96.3% for 64 of 69 segments in the collection, and it was not less than 46.5% for any segment, which is sufficient for segment and subtype identification. Nine isolates contained high priority H5, H7, and H9 segments, all of which had greater than 98.7% breadth of enrichment. This included two isolates from zoonotic human infections (H5N1 and H7N9), which were extensively enriched despite the absence of reference sequences from human infections in the target space used for probe design.

We further examined the five segments with less than 96.3% breadth of enrichment to understand why they were apparently not captured in full. First, we used the ProbeTools *capture* module to assess if the AIV_v1 panel lacked probes targeting their particular genome segment sequences. We observed that most positions without significant enriched were nonetheless extensively covered by the probe panel (Figure 5A). This indicated that insufficient design by ProbeTools was not a major explanation for the lack of significant capture of these segments.

**Figure 5:**
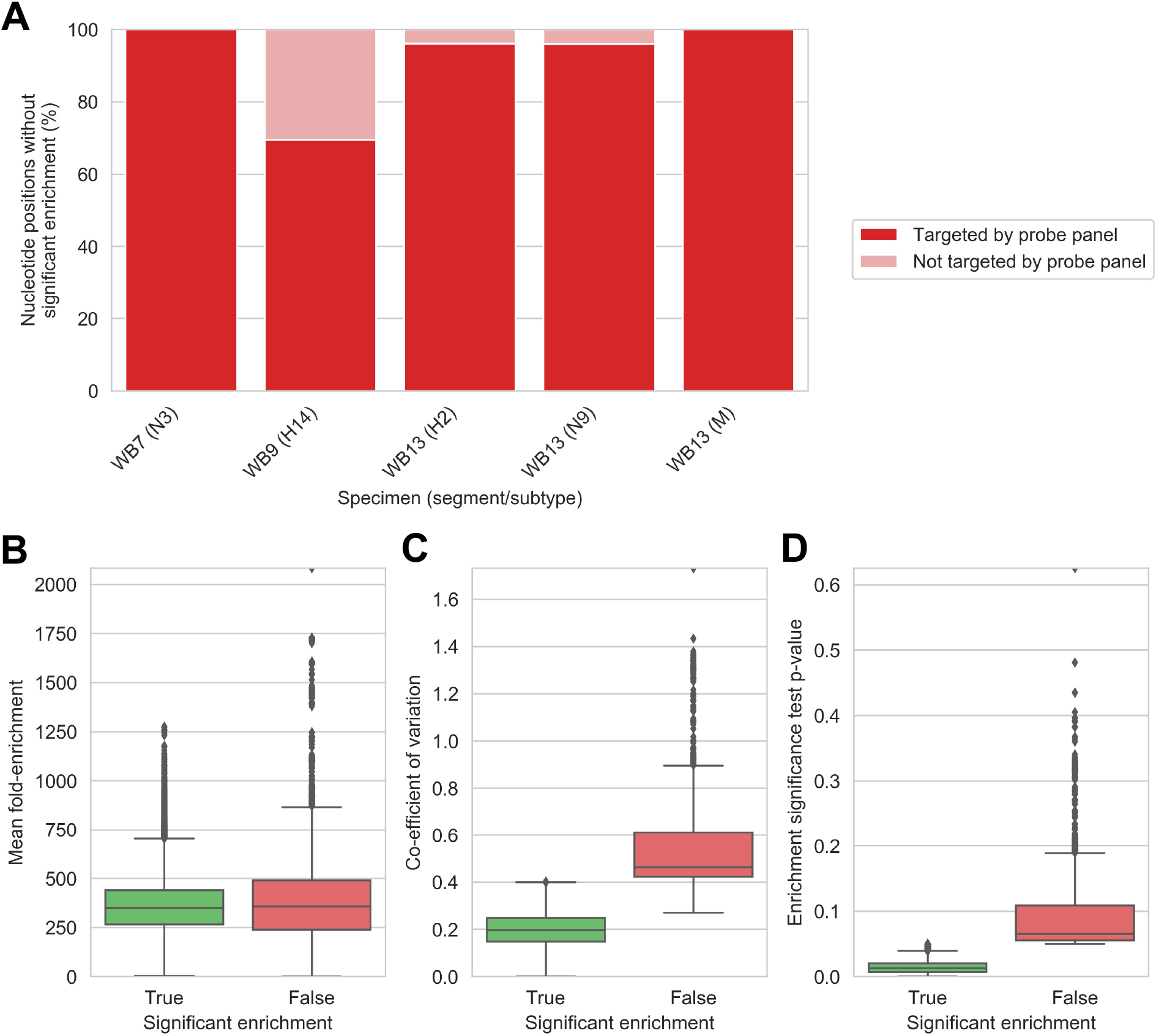
Lack of significant enrichment in segments with lower breadths of enrichment was due to experimental variation between capture replicates instead of insufficient probe design. A representative collection of 23 egg-cultured avian influenza viruses was captured three times independently using the ProbeTools-designed AIV_v1 panel. A) ProbeTools *capture* was used to predict probe panel coverage of positions without significant enrichment from 5 genome segments with breadths of enrichment less than 96%. These positions were extensively targeted by probes in the AIV_v1 panel. B) Fold-enrichment was comparable for positions with and without significant enrichment. The difference in distribution means was only 1.09-fold, although it was statistically significantly (p<0.0001, Welsh’s t-test) due to the large number of nucleotide positions involved in the comparison (n=96,376 and n=3,082 for positions with and without significant enrichment respectively). C) Variation in fold-enrichment between three independent replicates was significantly higher for positions that did not achieve significant enrichment (p<0.0001, Levene’s test). D) Most positions with insignificant enrichment narrowly failed the enrichment test’s pre-determined alpha level of 5%.

Next, we assessed whether experimental factors were responsible for nucleotide positions in these segments failing to achieve statistically significant enrichment. Fold-enrichment values between positions with and without significant enrichment were comparable, but variation between capture replicates were significantly different, with higher variation for positions that were not significantly enriched (Figure 5BC). Despite this source of experimental variation, and the limited number of replicates that was practical for us to perform, only 3.1% of nucleotide positions across all HA, NA, and M segments were impacted, and most of these positions only barely failed the enrichment significance test (half achieved a p-value < 0.07) (Figure 5D). Overall, our *in vitro* capture results demonstrated that the ProbeTools-designed AIV_v1 panel performed well on real viral isolates, effectively removing background material and providing high breadths of enrichment across HA, NA, and M segment targets.

### Comparison of *in silico* probe coverage prediction and *in vitro* probe capture enrichment

ProbeTools relies on *in silico* coverage assessment by the *capture* module, both for final panel evaluation and for identifying poorly covered sequences during incremental design. To validate ProbeTools’ coverage assessment algorithm, we examined how closely its *in silico* predictions agreed with *in vitro* capture results on egg-cultured AIV isolates.

Using the ProbeTools *capture* module, we determined which nucleotide positions in the egg-cultured AIVs were predicted to be covered by the AIV_v1 probe panel. We then compared these predictions to our *in vitro* capture results to see if significant enrichment had actually occurred at these nucleotide positions (Figure 6). Predicted probe coverage and significant enrichment results were concordant for 89.2% of nucleotide positions. Only 2.3% of nucleotide positions targeted by the AIV_v1 panel were not significantly enriched. These were concentrated in the five segments discussed above that were impacted by variability between replicates (Figure S1). We also noted that 7.7% of nucleotide positions were significantly enriched despite not being targeted by the AIV_v1 panel, a phenomenon that was observed in most segments across all isolates (Figure 6 and Figure S1). We attribute this to the capture of larger fragments containing untargeted sequences adjacent to the location annealed by the probe. It might also indicate that local alignment parameters used by ProbeTools *capture* are more conservative than actual annealing thermodynamics. Either way, these results showed that ProbeTools predictions generally reflected actual capture of target genomic material, and *in silico* predictions more often underestimated panel performance when predictions were incorrect.

**Figure 6:**
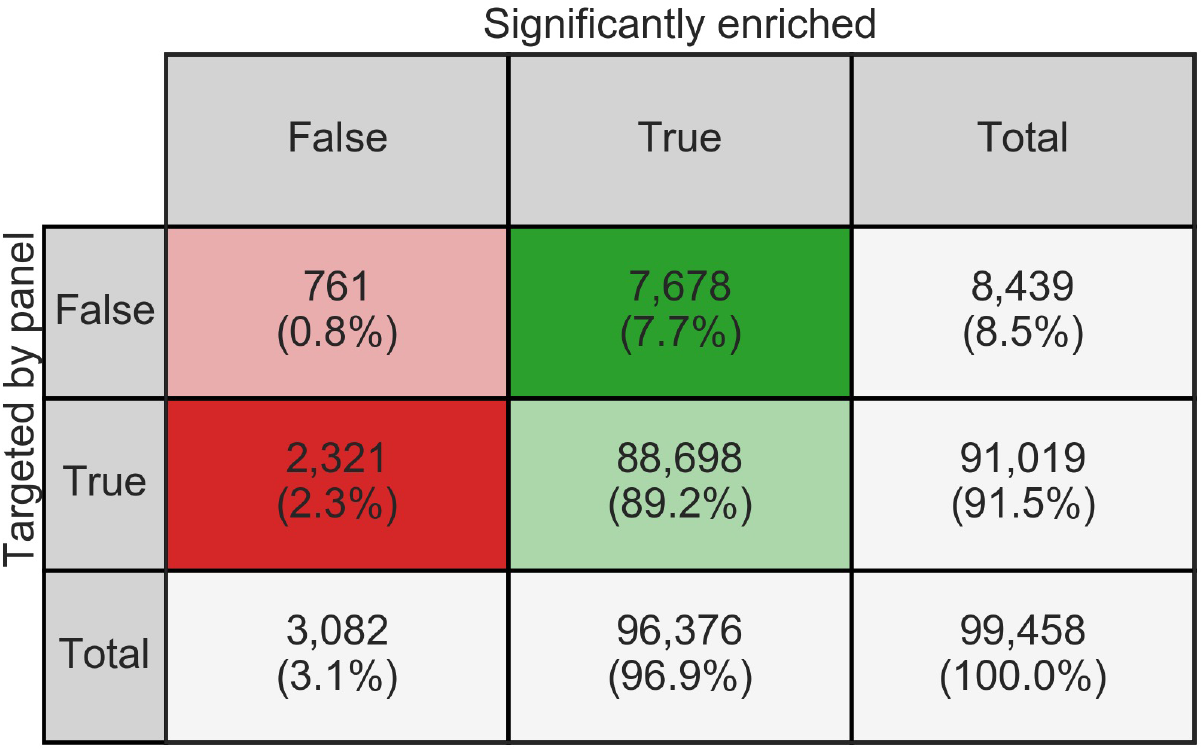
*In silico* predictions of probe coverage by ProbeTools were highly concordant with actual *in vitro* enrichment of egg-cultured AIV isolates. A representative collection of 23 egg-cultured avian influenza viruses was captured three times independently using the ProbeTools-designed AIV_v1 panel. Pre- and post-capture pools were sequenced to determine which nucleotide positions in the haemagglutinin (HA), neuramindase (NA), and matrix (M) genome segments of these isolates had been significantly enriched. The ProbeTools *capture* module was used to assess which nucleotide positions of these HA, NA, and M genome segments were targeted by the ProbeTools-designed panel. Each cell indicates the number of nucleotide positions meeting the corresponding *in silico* prediction and *in vitro* capture conditions.

## DISCUSSION

This study highlighted some important considerations when designing panels using ProbeTools. Foremost among these was the effect of target space composition on panel inclusivity. In this AIV case study, we noted a significant positive monotonic association between panel coverage and the number of reference sequences representing a particular subtype in the target space. Based on how the ProbeTools algorithm ranks probe candidates by the number of k-mers in the cluster they represent, it stands to reason that over-representing similar taxa (which would contain many similar k-mers) would bias the resulting panel towards these taxa.

Consequently, ProbeTools users should have a thorough knowledge of the contents of their target space and the possible sources of sampling bias in the databases from which they obtain their reference sequences. In the case of AIVs, the agricultural impacts and public health threats of certain HA subtypes have led to more frequent sequencing of these subtypes and accessioning of their genome sequences in popular databases. For our panel, this contributed to bias towards subtypes like H5, H7 and H9. Whether this is a benefit or limitation will depend on the intended application. In the context of outbreak prevention and pandemic preparedness, a panel biased towards taxa that are known for their agricultural impact and zoonotic potential is beneficial. If the objective is to characterize viral diversity and ecology in wildlife, however, this could be a limitation.

To obtain the best results, ProbeTools users should purposefully curate their target space to serve their probe capture objectives. Users may want to identify taxa whose detection is a priority and over-represent them in the target space. Conversely, users may want to ‘flatten’ their target space to ensure no particular taxa, clades, subtypes, *etc* dominate. This could be done manually, by selecting specific sequences to represent relevant groups, or it could be attempted bioinformatically by pre-clustering target sequences, providing the number and length of target sequences do not make this computationally prohibitive.

Another strategy could be to use the various ProbeTools modules to extract low coverage sequences from specific groups whose target sequences have poor probe coverage after a core panel is designed. For instance, had H15 subtype AIVs been a surveillance priority in this study, supplemental H15-specific probes could have been designed by running the *capture, getlowcov*, and *makeprobes* modules on the H15 subset of target sequences after noting their comparatively low coverage by the main panel. In this way, the modular nature of ProbeTools and the relatively simple-to-understand algorithms within each module empower users to experiment and find creative solutions. This flexibility is instrumental for tailoring probe panels to the needs of the user and their specific viral capture application.

## CONCLUSIONS

In this study, we used ProbeTools to create an effective and broadly inclusive panel of hybridization capture probes for subtyping AIVs. Our results show the utility of this panel as a tool for AIV surveillance, outbreak prevention, and pandemic preparedness. They also demonstrate that ProbeTools can effectively design probes against hypervariable genomic targets like avian-origin HA and NA segments. This validation of ProbeTools’ core design and coverage assessment algorithms shows that they are suitable for other challenging design applications, *e*.*g*. other viruses with hypervariable genes and pan-viral capture panels targeting multiple diverse taxa.

An increasing frequency of zoonotic outbreaks, epidemics, and pandemic crises has renewed interest in characterizing viral diversity at the interface of wildlife, livestock, game, and humans [39-42]. Genomic sequencing is becoming central to these One Health ventures, and viral capture panels will need designing and updating as our knowledge of viral threats continues to expand [43-44]. The on-going COVID-19 pandemic has also demonstrated the value of viral genomics to public health [45-48], resulting in unprecedented investments in sequencing capacity at public health laboratories. This will expand routine genomics for numerous common pathogens, requiring the development of new target enrichment protocols. ProbeTools can facilitate probe design tasks for all of these endeavours.

## METHODS

### ProbeTools modules

ProbeTools consists of five main modules written in Python (v3.7.3) that perform essential tasks in the probe design process. ProbeTools is freely available under the MIT License. It can be installed easily using the Anaconda/Miniconda package and environment manager. Alternatively, it can be installed via the Python Package Index, followed by separate installation of its VSEARCH and BLASTn dependencies. Installation instructions, source code, documentation, and usage examples are available at https://github.com/KevinKuchinski/ProbeTools.

The *<u>clusterkmers</u>* module enumerates and clusters probe-length k-mers from user-provided target sequences. 1) K-mers are enumerated using a sliding window that advances by a specified number of bases. 2) K-mers are clustered based on nucleotide sequence similarity using VSEARCH cluster_fast [34]. 3) Centroid sequences from each cluster are ranked by the size of the cluster they represent. Centroids from larger clusters are assumed to be better probe candidates by virtue of having similarity to more k-mers in the target space. By default, *clusterkmers* enumerates 120-mers, advancing the window one base at a time, and it clusters using a nucleotide sequence identity threshold of 90%. Previous studies have observed effective hybridization between targets and probes with this degree of sequence similarity [9, 11].

The *<u>capture</u>* module predicts how well user-provided probe sequences cover user-provided target sequences. 1) Each probe sequence is locally aligned against each target sequence using BLASTn [35]. 2) Alignments are filtered, retaining those with a minimum sequence identity over a minimum alignment length. 3) Subject alignment start and end coordinates are extracted from the BLASTn results to determine which nucleotide positions in the target sequences are covered by probes. By default, *capture* requires 90% sequence identity over at least 60 bases to assign probe coverage to the aligned positions.

The *<u>getlowcov</u>* module uses the output of *capture* to extract genomic regions with low coverage from the provided targets. This allows for additional probe design focused on poorly covered regions of the target space. This module returns all sub-sequences where a minimum number of consecutive bases were covered by fewer than a specified number of probes. By default, *getlowcov* returns all sub-sequences over 40 bases in length where all bases in the sub-sequence were covered by zero probes.

The *<u>stats</u>* module uses the output of *capture* to calculate coverage statistics. For each provided target, it calculates the percentage of nucleotide positions covered by varying numbers of probes (“target coverage” and “probe depth”).

The *<u>makeprobes</u>* module chains the previous modules together to implement a generalized incremental design strategy (Figure 2). In this strategy, probes are designed in batches, and regions of the target space with probe coverage are removed between batches so that additional probes are focused on poorly covered areas. This module can be used as a convenient departure point for custom designs. The user is only required to provide target sequences and select a batch size. They can optionally specify a maximum panel size and target space coverage goal. The *makeprobes* module iterates through its design loop, adding batches of probes to the panel until the maximum panel size is met, the target space coverage goal is achieved, or no further probes can be generated.

### Preparation of AIV target space

All available full-length influenza A virus genome segment sequences from avian hosts were downloaded from the Influenza Research Database (www.fludb.org) on Dec 5, 2017 [36]. Sequences containing degenerate bases were removed to avoid low quality entries. Sequences were then clustered using VSEARCH cluster_fast (v1.0.7) [34] with a 100% sequence identity threshold to remove redundant entries. The remaining entries were used as our final AIV target space (described in Table 2).

**Table 2:** Avian influenza virus reference sequences used as target space in this study. Full-length genome segment sequences from avian hosts were downloaded from the Influenza Research Database (www.fludb.org). Sequences containing degenerate bases were removed, then the remaining sequences were clustered using a 100% nucleotide sequence identity threshold to discard redundant entries. This provided a final target space of 36,313 reference sequences representing all avian-origin haemagglutinin (HA) subtypes, neuraminidase (NA) subtypes, and matrix (M) segments.

| Genome segment | Subtype | Reference sequences in target space (#) | Target space size (KB) |
| --- | --- | --- | --- |
| HA | H1 | 539 | 939.1 |
|  | H2 | 381 | 664.0 |
|  | H3 | 1,309 | 2,267.6 |
|  | H4 | 1,135 | 1,944.1 |
|  | H5 | 3,546 | 6,129.7 |
|  | H6 | 1,378 | 2,361.3 |
|  | H7 | 1,266 | 2,148.5 |
|  | H8 | 122 | 209.9 |
|  | H9 | 2,653 | 4,498.9 |
|  | H10 | 591 | 1,005.4 |
|  | H11 | 520 | 897.5 |
|  | H12 | 176 | 301.5 |
|  | H13 | 232 | 405.4 |
|  | H14 | 22 | 38.3 |
|  | H15 | 13 | 22.7 |
|  | H16 | 156 | 271.7 |
|  | HA untyped | 733 | 1,254.8 |
|  | **HA total** | **14,772** | **25,360.4** |
| NA | N1 | 2,742 | 3,804.9 |
|  | N2 | 3,162 | 4,498.5 |
|  | N3 | 945 | 1,347.3 |
|  | N4 | 174 | 249.8 |
|  | N5 | 296 | 427.4 |
|  | N6 | 1,424 | 2,037.0 |
|  | N7 | 502 | 718.4 |
|  | N8 | 1,272 | 1,822.9 |
|  | N9 | 667 | 948.5 |
|  | NA untyped | 783 | 1,116.6 |
|  | **NA total** | **11,967** | **16,971.0** |
| **M** | **none** | **9,574** | **9,582.4** |

### AIV_v1 probe panel design

The AIV_v1 panel was designed against our final AIV target space using the ProbeTools *makeprobes* module as follows: 2,000 probes were designed against HA targets in 20 batches of 100 probes; 1,500 probes were designed against NA targets in 15 batches of 100 probes, and 200 probes were designed against M targets in 20 batches of 10 probes. All designs were conducted using *makeprobes*’s default parameters with ProbeTools v0.0.5, VSEARCH v1.0.7, and BLASTn v2.2.31.

The top-ranked 1,935 HA probes, 1,435 NA probes, and 184 M probes were combined into the final panel. Additional probes were added to the panel for potential control and validation applications, including 36 probes targeting the common reference strain A/Puerto Rico/8/34 and 10 probes targeting synthetic spike-in DNA oligomers with randomly generated artificial sequences. This provided a final panel of 3,600 probes (a breakpoint in the manufacturer’s pricing structure), which was synthesized as a custom panel by Twist Bioscience (San Francisco, CA, USA). Sequences for probes in the AIV_v1 panel are provided in Supplemental Material 1.

### Preparation of sequencing libraries from egg-cultured influenza isolates

Detailed laboratory procedures for the following are provided in Supplemental Material 2. RNA extracted from egg-cultured AIV isolates was provided by the Canadian Food Inspection Agency’s National Centre for Foreign Animal Disease (Winnipeg, Manitoba, Canada) and the Public Health Agency of Canada’s National Microbiology Laboratory (Winnipeg, Manitoba, Canada). cDNA was prepared from each isolate using a previously described method [37]. cDNA was also prepared from a mock-infected egg culture to generate background genomic material for diluting capture pools. cDNA was fragmented by sonication, then prepared into sequencing libraries for Illumina platforms with unique dual index barcodes. Adapter-ligated cDNA was split into three separate barcoding reactions, providing three separately barcoded replicate libraries for each isolate.

### Probe capture enrichment and genomic sequencing of libraries prepared from egg-cultured influenza isolates

Detailed laboratory and bioinformatic procedures for the following are provided in Supplemental Material 2. 1) Three pools were prepared, with each pool containing one replicate library from each AIV isolate. These pools were sequenced in-house on Illumina MiSeq to generate full HA, NA, and M segment sequences for each isolate and to confirm HA and NA subtypes. 2) Each pool was diluted in 1:100 (ng/ng) in one of three replicate libraries of background genomic material that had been prepared from a mock-infected chicken egg. Aliquots of each diluted pool were sequenced pre-capture at Canada’s Michael Smith Genome Sciences Centre (Vancouver, BC) on one Illumina HiSeq X lane to establish baseline HA, NA, and M segment abundance. 3) Each diluted pool was independently captured using the AIV_v1 probe panel. Captured pools were then sequenced in-house on Illumina MiSeq to assess target enrichment of HA, NA, and M segments post-capture.

### Analysis of significant probe capture enrichment for egg-cultured AIV isolates

1) Pre- and post-capture depths of coverage were determined by mapping each library’s sequencing reads to the HA, NA, and M segment sequences of its corresponding AIV isolate. 2) Depths of coverage were normalized by dividing raw pre- and post-capture read depths by the total reads in the corresponding pre- and post-capture pools (Table S2). 3) For each library, fold-enrichment at each nucleotide position was calculated by dividing the normalized post-capture read depth by the normalized pre-capture read depth. 4) For each AIV isolate, mean fold-enrichment was calculated at every nucleotide position from the fold-enrichment values of its three independently captured replicate libraries. 5) Mean fold-enrichment values and their standard deviations were used to determine if significant enrichment had occurred at all nucleotide positions using a one-sample T-test against the fixed value of one-fold enrichment with an alpha level of 5%.

## DECLARATIONS

### Ethics approval and consent to participate

Not applicable.

### Consent for publication

Not applicable.

### Availability of data and materials

ProbeTools v0.0.5 source code, which was used to design the final probe panel and assess its coverage of target sequences *in silico* for this manuscript, is available on GitHub at https://github.com/KevinKuchinski/ProbeTools. FASTA files of the HA, NA, and M genome segment reference sequences used as a target space for design and assessment in this manuscript (described in Table 2) are provided as part of the ProbeTools v0.0.5 release. The sequences of the AIV_v1 probe panel are also provided as part of the ProbeTools v0.0.5 release, and they are also included in this manuscript’s supplemental information as Supplemental Material 1. Data from the *in vitro* captures are provided in BAM format with pre- and post-capture libraries mapped to the HA, NA, and M genome segment sequences of their corresponding egg-cultured AIV isolate. These can be accessed from the NCBI Short Read Archive as part of BioProject PRJNA796698. Total read counts used to normalize depths of coverage in these libraries are provided in the manuscript’s supplemental material as Table S2.

### Authors’ contributions

KK designed and implemented the ProbeTools algorithms, wrote the ProbeTools source code, designed the AIV_v1 probe panel, prepared sequencing libraries, performed probe captures and in-house sequencing, analyzed the data, and wrote the manuscript. JD performed preliminary studies with k-mer clustering, assisted with the design and implementation of the ProbeTools algorithms, and provided guidance on bioinformatic data analysis. CH helped assemble the validation collection of egg-cultured AIV isolates, ensured relevant strains were included, and provided direction for AIV probe panel design to ensure its suitability for agricultural surveillance applications. WH provided guidance on implementing ProbeTools algorithms, best practices for constructing and distributing bioinformatics tools and packages, and bioinformatic data analysis. NP provided guidance on experiment design for *in vitro* captures, troubleshooting for library preparation, probe capture, and sequencing of egg-cultured AIV isolates, and provided direction for AIV probe panel design to ensure its suitability for public health surveillance applications. All authors reviewed and contributed comments on the manuscript.

## Acknowledgements

We would like to acknowledge the efforts of all laboratories world-wide who have submitted genomic sequences to the Influenza Research Database. Dr. Yohannes Berhane and Matthew Suderman at the Canadian Food Inspection Agency’s National Centre for Animal Disease were instrumental in providing diverse egg-cultured AIV validation material from wild birds and poultry. We also thank Dr. Agatha Jassem at the British Columbia Centre for Disease Control’s Public Health Laboratory and Dr. Nathalie Bastien at the Public Health Agency of Canada’s National Microbiology Laboratory for providing H5N1 and H7N9 validation material from human infections. Additionally, we thank Tracy Lee at the British Columbia Centre for Disease Control’s Public Health Laboratory for providing primers used to generate cDNA from AIV egg-cultures.

