## Supplemental Material S2 for "ProbeTools: Designing hybridization probes for targeted genomic sequencing of diverse and hypervariable viral taxa"

### MATERIALS AND METHODS SUPPLEMENTAL

**Preparation of cDNA from egg-cultured AIV isolates and mock-infected egg culture background material:** cDNA was prepared from RNA extracts using the method by Zhou *et al.* [1]. 20 µl RT-PCR reactions were prepared using the Invitrogen SuperScript IV One-Step RT-PCR System (#12594025) with 5 µl of input RNA. Primers MBTuni-12 (5'-ACGCGTGATCAGCAAAAGCAGG-3') and MBTuni-13 (5'-ACGCGTGATCAGTAGAAACAAGG-3') were both added at a final concentration of 4 µM. The following PCR cycling conditions were used: 1 cycle of 42 °C for 30 mins, then 98 °C for 30 secs; 5 cycles of 98 °C for 30 secs, then 45 °C for 45 secs, then 72 °C for 3 mins; 45 cycles of 98 °C for 30 secs, then 60 °C for 45 secs, then 72 °C for 3 mins; 1 cycle of 72 °C for 10 mins. cDNA products were purified using 1X SPRI beads from NEBNext Ultra II DNA Library Prep with Sample Purification Beads (#E7103). Bead clean-up reactions were washed twice with 200 µl of 80% ethanol and eluted in 20 µl of NEBNext 0.1X TE buffer.

**Construction of sequencing libraries from cDNA:** cDNA was fragmented using the Covaris ME220 instrument. 5 µl of purified cDNA was added to 50 µl of nuclease-free water in Covaris 8 microTUBE-50 AFA FiberStrip V2 tubes (#520174). The following shearing conditions were used: 20 °C water temperature, 50 W peak power, 20% duty factor, 1000 cycles per burst, and 75 sec duration.

50 ul of fragmented cDNA was used as input for the NEBNext Ultra II End Repair/dA-tailing module (#E7546) with reactions prepared according to the manufacturer's protocol and incubated at 20 °C for 30 mins then 65 °C for 30 mins.

Afterwards, these reactions were directly used as input for the NEBNext Ultra II Ligation module (#E7595) with reactions prepared according to the manufacturer's protocol and incubated at 20 °C for 30 mins. A USER digest was then performed as directed by the manufacturer's protocol with a 30 min incubation at 37 °C. Adapter-ligated cDNA was purified using 0.8X SPRI beads from NEBNext Ultra II DNA Library Prep with Sample Purification Beads (#E7103). Bead clean-up reactions were washed twice with 200 µl of 80% ethanol and eluted in 50 µl of NEBNext 0.1X TE buffer.

Purified adapter-ligated cDNA from each specimen was aliquoted into three separate library barcoding PCRs, creating three replicate libraries for each specimen. Barcoding PCRs were conducted using the NEBNext Multiplex Oligos for Illumina (96 Unique Dual Index Primer Pairs) kit (#6440). Barcoding PCRs used the following cycling conditions: 1 cycle of 98 °C for 1 min; 10 cycles of 98 °C for 30 secs, then then 65 °C for 75 secs; 1 cycle of 65 °C for 10 mins. Barcoding PCRs were purified using 0.8X SPRI beads from NEBNext Ultra II DNA Library Prep with Sample Purification Beads (#E7103). Bead clean-up reactions were washed twice with 200 µl of 80% ethanol and eluted in 20 µl of NEBNext 0.1X TE buffer.

**Creation of replicate capture pools and their enrichment by hybridization probe capture:** Replicate library pools were constructed by combining 100 ng of one replicate library from each specimen. These pools were sequenced in-house on Illumina MiSeq with a V3 (600 cycle) reagent kit (MS-102-3003) to generate HA, NA, and M segment sequences (see below). 12 ng of each pool was diluted in 1188 ng of background material from one of three libraries generated from mock-infected egg cultures (1:100 ng/ng dilution). 750 ng from each diluted libraries pool was used as capture pools. Each pool was captured independently on different days.

Prior to hybridization, capture pools were completely evaporated in a vacuum oven at 50 °C and -20 mm Hg. Dried pools were used to set up hybridization reactions with 0.2 fmol/probe of our custom avian influenza probe panel (Twist Biosciences, San Francisco, CA, USA), Twist Universal Blockers (#100578), and the Twist Hybridization Reagents kit (#100930) following the manufacturer's protocol. Hybridization reactions were incubated at 70 °C for 16 hours, then washed with the Twist Wash Buffers kit (#100985) following the manufacturer's protocol until the final step, at which point the streptavidin bead slurry was resuspended in 22.5 µl of nuclease-free water instead of 50 µl. The entire 22.5 µl volume was used in the post-capture PCR, which was set up with NEBNext Ultra II Q5 2X Master Mix (#M0544), and Illumina amplification primers from the Twist Hybridization Reagents kit (#100930). Post-capture PCRs were conducted with the following cycling conditions: 1 cycle of 98 °C for 45 secs; 10 cycles of 98 °C for 15 secs, then then 60 °C for 30 secs, then 65 °C for 30 secs; 1 cycle of 65 °C for 10 mins. Post-capture PCRs were purified using 0.8X SPRI beads from NEBNext Ultra II DNA Library Prep with Sample Purification Beads (#E7103). Bead clean-up reactions were washed twice with 200 µl of 80% ethanol and eluted in 20 µl of NEBNext 0.1X TE buffer.

**Generation of HA, NA, and M segment sequences for egg-cultured AIV isolates:** Before processing, FASTQ files from each isolate's three replicate libraries were concatenated together. Low-quality bases were trimmed from read sequences using sickle pe (v1.33) with default parameters. Illumina adapter sequences were trimmed from read sequences using cutadapt (v3.3) with default parameters based on the sequence 5'-AGATCGGAAGAGC-3'. Cutadapt (v3.3) was also used to trim sequences for cDNA synthesis primers MBTuni12 and MBTuni13.

Trimmed reads were assembled *de novo* into contigs using SPAdes (v3.15.3) with the rnaviral and isolate options selected. Contigs were locally aligned against avian influenza A virus reference sequences with BLASTn (2.12.0) to identify those representing HA, NA, and M segments and to ensure their completeness. Trimmed reads were then mapped to the HA, NA, and M segment contigs using bwa mem (0.7.17) with default parameters. Output alignments files were filtered using samtools view (v1.14) using parameters -f 3, -F 2828, and -q 30 to discard unmapped reads, reads with low mapping scores, reads without properly mapped mates, and non-primary mappings. Filtered alignment files were then sorted with samtools sort (v1.14) and indexed with samtools index (v1.14). Variants were called from filtered alignment files using bcftools mpileup and call (v1.14). Bcftools mpileup required minimum read and mapping qualities of 30 and a minimum depth of coverage of 5, while bcftools call used a ploidy of 1. Variants were applied to reference sequences using bcftools consensus (v1.14), making positions supported by fewer than 5 reads with the N ambiguity character. Leading and trailing runs of Ns were then trimmed to produce final HA, NA, and M segment sequences.

**Analyzing depth of coverage for pre-capture and post-capture libraries:**

Reads from pre- and post-capture libraries were mapped to their corresponding HA, NA, and M segment sequences (as generated above) using bwa mem (v0.7.17). Output alignment files were filtered using samtools view (v1.14) with parameters -f 3, -F 2828, and -q 30 to discard unmapped reads, reads with low mapping scores, reads without properly mapped mates, and non-primary mappings. Filtered alignment files were then sorted with samtools sort (v1.14) and indexed with samtools index (v1.14).

Depth of coverage pre- and post-capture was determined for each library with bedtools genomecov (v2.30.0). Depth of coverage tables in TSV format were loaded into Pandas (v0.24.2) dataframe objects in Python (v3.7.3). One-sample T tests were conducted using the ttest_1samp function in the SciPy package (v1.3.0).
